# Evaluation of the Ultima Genomics UG 100™ sequencing platform for clinical services

**DOI:** 10.64898/2025.12.11.693628

**Authors:** Luca Santuari, Ilya Kolpakov, Anthony Blin, Ioannis Xenarios, Cédric Howald

**Affiliations:** Health 2030 Genome Center, Chem. des Mines 9, Geneva, 1202, Switzerland; University of Lausanne, Quartier Centre, Lausanne, 1015, Switzerland

**Keywords:** ultima genomics, whole-genome sequencing, small variant calling, clinical accreditation

## Abstract

The steep reduction in the cost of genome sequencing started with the introduction of Solexa Sequencing-By-Synthesis technology in 2006 has plateaued recently due to technical limitations in the use of closed flowcells and to the cost of reagents. The Ultima Genomics UG 100™ is the first sequencing machine to lower the cost of human genome sequencing to 80$. However, technical limitations in resolving long homopolymer regions undermine the application of this technology to short variant calling in clinical settings. Here, we evaluate the ability of UG 100™ to identify short variants in relation to its suitability for clinical accreditation, by comparing it with the Illumina NovaSeq 6000 Systems platform. We focus specifically on the small variant calling performance in long homopolymer regions, both genome-wide and in relation to a set of medically-relevant genes that are challenging to sequence. Our analysis aims at supporting clinicians in determining whether the UG 100™ platform is well-suited for their studies, and to guide clinical sequencing centers in evaluating the adoption of this emerging technology.

## Introduction

From the start of the Human Genome Project in 1990, sequencing a human genome has dropped from an estimate of 3B$ to below 1000$ [22]. However, recently sequencing costs have stalled due to technical limitations in the use of closed flowcells and to the cost of sequencing reagents. In an effort to further reduce sequencing costs, in February 2024 Ultima Genomics released the UG 100™ platform, a high-throughput sequencing device that lowers the cost of human genome sequencing to 80$. This is achieved by the use of an open flowcell and an efficient use of reagents [1].

The sequencing approach the UG 100™ instrument adopts is called mostly natural Sequencing-By-Synthesis (mnSBS) [1]. The device uses an open wafer disc with a diameter of 200 mm, where reagents are delivered by nozzles near the center of the disc and distributed by centrifugal force. At each sequencing cycle, a single base is added to the wafer in a mixture of non-terminated fluorescently labelled (< 20%) and unlabeled nucleotides. At this step, zero, one or more nucleotides are incorporated into the growing strand of the synthesized DNA and the fluorescent signal is read by optical scanning. This is similar to the process of reading a compact disc. A beneficial property of this approach is that the fluorescent signal is proportional to the length of the homopolymer that is being resolved on the DNA template, since it is proportional to the number of incorporated fluorescently labelled nucleotides.

However, the accuracy of homopolymer calling drops below 90% after a length of 8 bases ([1], Fig.1E and G).

**Fig. 1.**
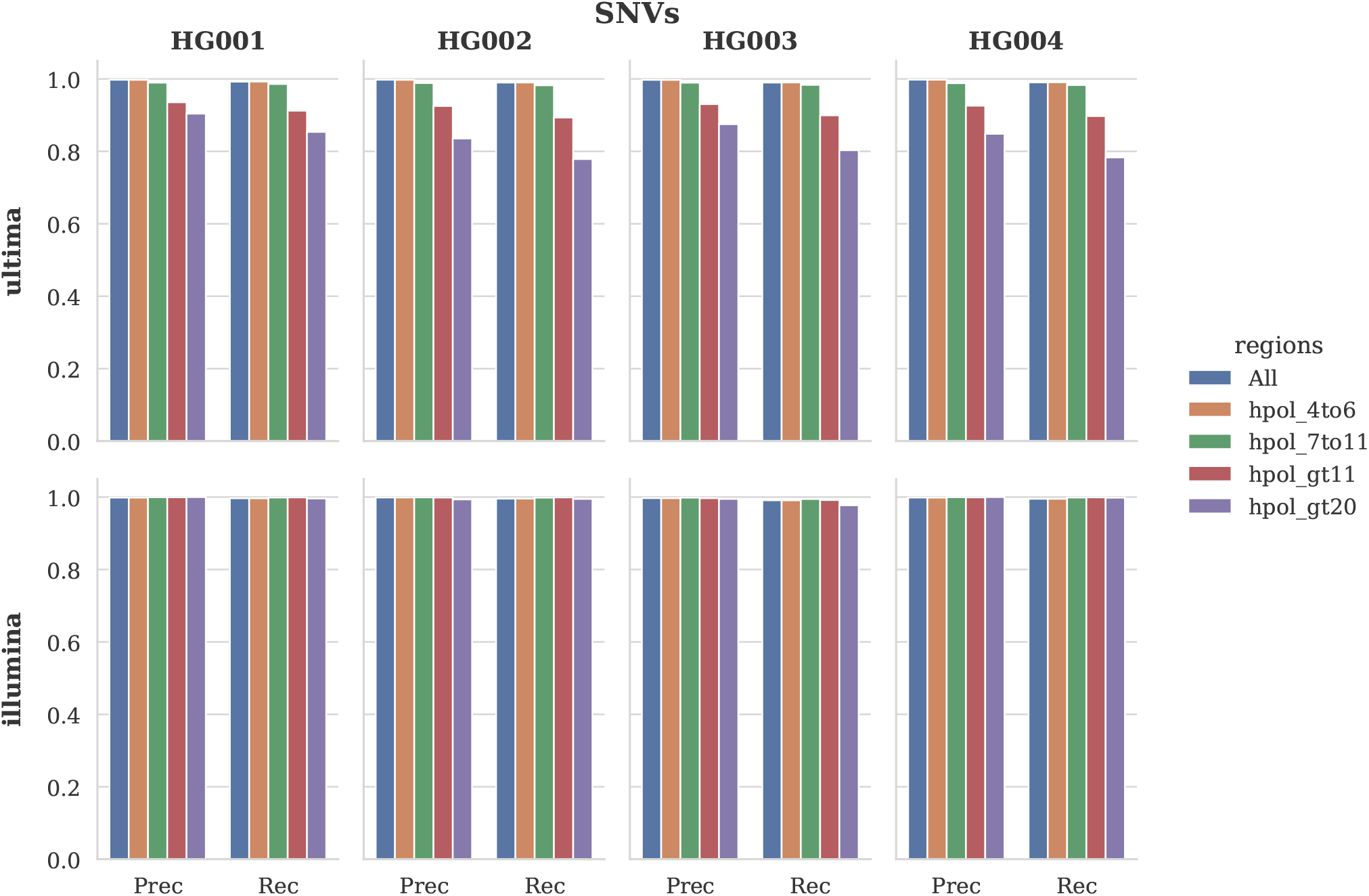
Precision (Prec) and recall (Rec) for SNV calling on the four GIAB samples sequenced with UG 100™ (ultima) and NovaSeq 6000 System (illumina). Genomic regions considered are all benchmark regions (All), homopolymers from 4 to 6 bp (hpol_4to6), homopolymers from 7 to 11 bp (hpol_7to11), homopolymers longer than 11 bp (hpol_gt11) and homopolymers longer than 20 bp (hpol_gt20).

Currently the sequencing machines of choice for clinical applications in terms of cost-quality and widespread adoption are the Illumina instruments of the NovaSeq series (NovaSeq 6000 System and NovaSeq X Series). The introduction of the Ultima Genomics UG 100™ platform in clinical settings holds the promise to further reduce the cost of sequencing a patient’s genome and opens up the possibility of scaling clinical studies. Accreditation of sequencing centers for clinical services requires a rigorous evaluation of the ability of the selected sequencing instrument to correctly identify the full extent of genomic variation in a patient’s genome, particularly with respect to clinically relevant regions. To this purpose, the Genome in a Bottle (GIAB) Consortium of the National Institute of Standards and Technology (NIST)’s Joint Initiative for Metrology in Biology (JIMB) has created a benchmark dataset with a gold standard set of high quality small variants (SNVs and Indels) for several well-characterized human genomes, identified using multiple orthogonal sequencing technologies and variant calling algorithms [19].

In this study, we evaluate the ability of the UG 100™ instrument to identify small variants in four well-characterized human genomes (HG001, HG002, HG003 and HG004), in comparison to the Illumina NovaSeq 6000 System, with a particular focus to long homopolymer regions, that are excluded from the High Confidence Regions (HCR) genomic tracks Ultima Genomics provides for the analysis [1]. To further evaluate the UG 100™ platform for clinical applications, we assess the small variant calling performance in a set of challenging medically-relevant genes for which a benchmark dataset is available [20].

### Evaluation of UG 100™’s short variant calling performance using the GIAB benchmark

We sought to assess the performance of short variant calling pipelines using the Ultima Genomics (UG) UG 100™ device with the UG VariantCalling workflow (ultima) [6] in comparison with the Illumina NovaSeq 6000 System processed with the DRAGEN v4.3.13 workflow (illumina). In a pilot study, we sequenced four genomes with broad consent: the pilot genome from the HapMap project NA12878 (NIST ID: HG001) and a trio of Ashkenazi Jewish ancestry from the Personal Genome Project: NA24385 (son, NIST ID:HG002), NA24149 (father, NIST ID:HG003) and NA24143 (mother, NIST ID: HG004) [24]. These genomes are part of the GIAB NIST v.4.2.1 small variant benchmark [19], a curated dataset of SNVs and Indels that have been established based on a consensus of orthogonal sequencing technologies and variant calling algorithms. This benchmark enables rigorous evaluation of sequencing pipelines for clinical applications.

The Global Alliance for Genomics and Health (GA4GH) has defined a set of best practices for benchmarking germline small variants [11]. Hap.py [11, 9] is a tool developed by Illumina that adheres to these best practices. It allows to calculate the precision and recall of a small variant callset (query) with respect to a gold standard callset (truth). In particular, hap.py preprocessing step takes into account that the same variant can have multiple different representations in the VCF file format [11].

Sequencing reads were generated with the Illumina NovaSeq 6000 and with the Ultima Genomics UG 100™ platforms. Reads were mapped on the GRCh38 reference genome. We used hap.py with the truth sets and confident regions from NIST v.4.2.1 to calculate the precision (TP/(TP+FP)) and recall (TP/(TP+FN)) for the small variant callsets identified in the four GIAB genomes using the ultima and illumina pipeline. In particular, to evaluate the performance in homopolymer regions, we used the stratification files generated by the GA4GH (v3.6) [3]. These regions include an extension of 5 bp in both directions. We considered different homopolymer length ranges: from 4 to 6 bp (hpol 4to6), from 7 to 11 bp (hpol 7to11), longer than 11 bp (hpol gt11), and longer than 20 bp (hpol gt20).

For SNVs (Fig.1), the precision and recall of ultima is lower than illumina in homopolymers longer than 11 bp (ultima precision: 0.929 ± 0.004, recall: 0.9 ± 0.008; illumina precision: 0.998 ± 0.001, recall: 0.997 ± 0.003). For homopolymers longer than 20 bp, both precision and recall of ultima further drop with respect to illumina (ultima precision: 0.865 ± 0.03, recall: 0.804 ± 0.034; illumina precision: 0.997 ± 0.003, recall: 0.991 ± 0.009). However, it is for Indels that ultima shows the largest decrease in performance with respect to illumina in homopolymers longer than 11 bp (ultima precision: 0.594 ± 0.019, recall: 0.507 ± 0.023; illumina precision: 0.996 ± 0.003, recall: 0.994 ± 0.006) and in homopolymers longer than 20 bp (ultima precision: 0.382 ± 0.018, recall: 0.136 ± 0.02; illumina precision: 0.982 ± 0.013, recall: 0.981 ± 0.019). These results are worse than those obtained using the publicly available Ultima Genomics GIAB dataset that has been generated with the newest Solaris chemistry (Fig.4). This chemistry improves both precision and recall values for HG001, HG002, HG003 and HG004, in particular in homopolymers longer than 11 bp for SNVs (precision: 0.955 ± 0.000, recall: 0.955 ± 0.002) and especially for Indels (precision: 0.866 ± 0.001, recall: 0.716 ± 0.017) (Fig.5). This accounts for an improvement in Indel calling of 45.8% in precision and 41.2% in recall. It is worth noting that for the NA24631 (NIST ID: HG005) genome, the son from the GIAB trio of Chinese ancestry, which is not included in our study, the Indels calling performance is better than the other four GIAB genomes, both in homopolymers longer than 11 bp (precision: 0.927 ± 0.0, recall: 0.867 ± 0.002) and in homopolymers longer than 20 bp (precision: 0.846 ± 0.009, recall: 0.860 ± 0.03).

### ClinVar pathogenic small variants in long homopolymer regions

ClinVar [12] is a public archive of the National Center for Biotechnology Information (NCBI) that contains germline and somatic variants annotated with clinical significance information. This represents a valuable resource to evaluate the extent of genomic variation for a certain clinical significance category that is present in regions challenging to sequence. We calculated the number and proportion of SNVs and Indels present in the ClinVar database (clinvar 20251123) that are overlapping with homopolymer regions longer than 11 bp, which are currently difficult to resolve with the ultima pipeline. Out of a total of 82,029 Pathogenic SNVs, only 69 overlap with homopolymers longer than 11 bp, and only 16 with homopolymers longer than 20 bp (Table 1). For Pathogenic Indels, out of a total of 91,686 variants, only 35 overlap with homopolymers longer than 11 bp, and only 9 with homopolymers longer than 20 bp (Table 2). This analysis is restricted to small variants with a single clinical significance (CLNSIG) term reported per variant using the standard terminology as recommended by the American Collect of Medical Genetics and Genomics (ACMG), the Association for Molecular Pathology (AMP) and the College of American Pathologists (CAP) [16]: benign, likely benign, uncertain significance, likely pathogenic and pathogenic. Several variants of uncertain significance (VUS) can be found in homopolymers longer than 11 bp (758 SNVs, 1184 Indels) and homopolymer longer than 20 bp (160 SNVs, 342 Indels).

**Table 1.** Overlap of ClinVar SNVs with homopolymer regions of varying lengths (in base pairs, bp), stratified by CLNSIG clinical significance categories. For each CLNSIG category, the percentage of overlap represents the proportion of ClinVar SNVs that fall within homopolymer regions relative to the total number of SNVs in that category.

| HPol (bp) | CLNSIG | Overlap | Total | % |
| --- | --- | --- | --- | --- |
| 4-6 | Benign | 48677 | 180246 | 0.2701 |
|  | Likely_benign | 261084 | 971105 | 0.2689 |
|  | Likely_pathogenic | 15228 | 70856 | 0.2149 |
|  | Pathogenic | 16696 | 82029 | 0.2035 |
|  | Uncertain_significance | 458592 | 2059317 | 0.2227 |
| 7-11 | Benign | 3116 | 180246 | 0.0173 |
|  | Likely_benign | 12259 | 971105 | 0.0126 |
|  | Likely_pathogenic | 435 | 70856 | 0.0061 |
|  | Pathogenic | 377 | 82029 | 0.0046 |
|  | Uncertain_significance | 10981 | 2059317 | 0.0053 |
| > 11 | Benign | 1245 | 180246 | 0.0069 |
|  | Likely_benign | 2201 | 971105 | 0.0023 |
|  | Likely_pathogenic | 86 | 70856 | 0.0012 |
|  | Pathogenic | 69 | 82029 | 0.0008 |
|  | Uncertain_significance | 758 | 2059317 | 0.0004 |
| > 20 | Benign | 291 | 180246 | 0.0016 |
|  | Likely_benign | 401 | 971105 | 0.0004 |
|  | Likely_pathogenic | 10 | 70856 | 0.0001 |
|  | Pathogenic | 16 | 82029 | 0.0002 |
|  | Uncertain_significance | 160 | 2059317 | 0.0001 |

**Table 2.**
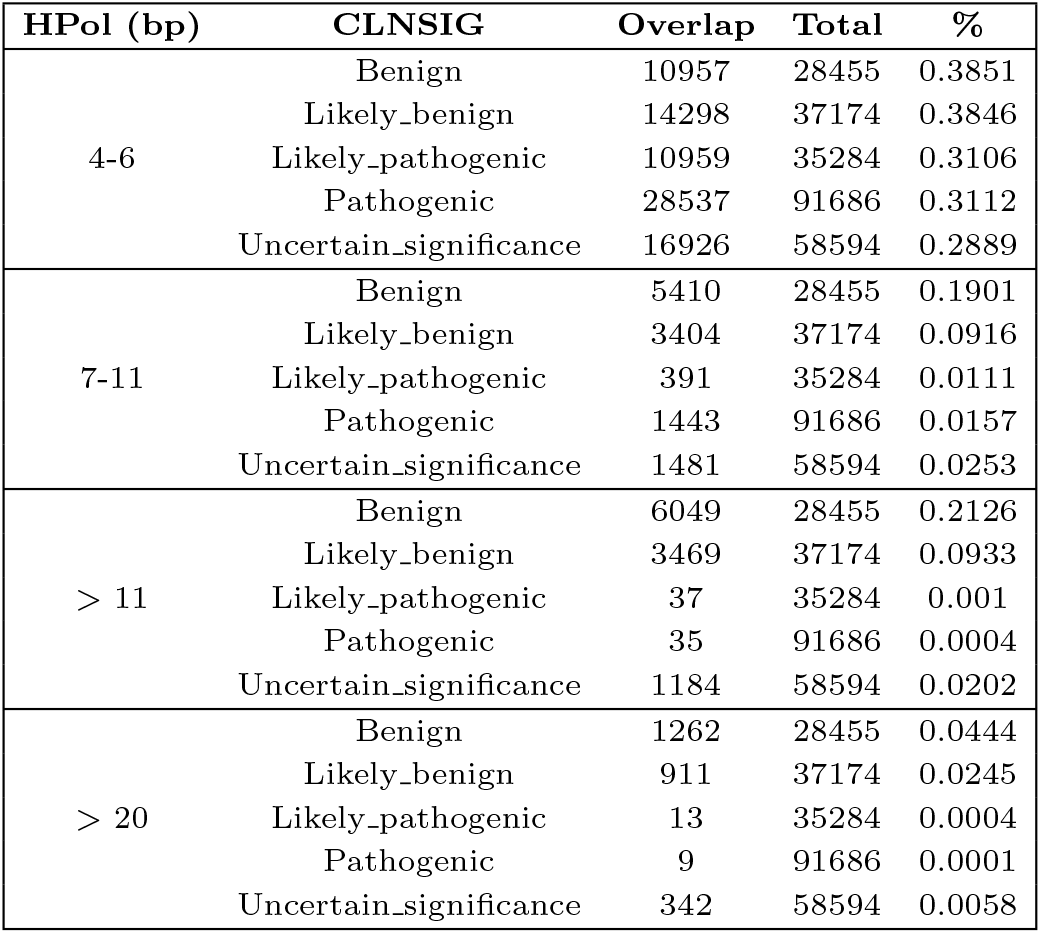
Overlap of ClinVar Indels with homopolymer regions of varying lengths (in base pairs, bp), stratified by CLNSIG clinical significance categories. For each CLNSIG category, the percentage of overlap represents the proportion of ClinVar Indels that fall within homopolymer regions relative to the total number of Indels in that category.

| HPol (bp) | CLNSIG | Overlap | Total | % |
| --- | --- | --- | --- | --- |
| 4-6 | Benign | 10957 | 28455 | 0.3851 |
|  | Likely_benign | 14298 | 37174 | 0.3846 |
|  | Likely_pathogenic | 10959 | 35284 | 0.3106 |
|  | Pathogenic | 28537 | 91686 | 0.3112 |
|  | Uncertain_significance | 16926 | 58594 | 0.2889 |
| 7-11 | Benign | 5410 | 28455 | 0.1901 |
|  | Likely_benign | 3404 | 37174 | 0.0916 |
|  | Likely_pathogenic | 391 | 35284 | 0.0111 |
|  | Pathogenic | 1443 | 91686 | 0.0157 |
|  | Uncertain_significance | 1481 | 58594 | 0.0253 |
| > 11 | Benign | 6049 | 28455 | 0.2126 |
|  | Likely_benign | 3469 | 37174 | 0.0933 |
|  | Likely_pathogenic | 37 | 35284 | 0.001 |
|  | Pathogenic | 35 | 91686 | 0.0004 |
|  | Uncertain_significance | 1184 | 58594 | 0.0202 |
| > 20 | Benign | 1262 | 28455 | 0.0444 |
|  | Likely_benign | 911 | 37174 | 0.0245 |
|  | Likely_pathogenic | 13 | 35284 | 0.0004 |
|  | Pathogenic | 9 | 91686 | 0.0001 |
|  | Uncertain_significance | 342 | 58594 | 0.0058 |

### Performance on the CMRG benchmark

The GIAB benchmark excludes a set of challenging medically-relevant genes due to their repetitive content and sequence complexity. To overcome this, Wagner et al. [20] curated a small variant benchmark for the HG002 genome for a set of 273 challenging medically-relevant genes (CMRG). We evaluated the variant calling performance of the ultima pipeline with respect to the illumina pipeline on this benchmark, with a particular focus to long homopolymer regions, by using the GA4GH stratifications. Although perfect and imperfect homopolymers longer than 20 bp, plus 5 bp on each side, were excluded from the benchmark, this variant set enables evaluation of the ultima dataset for homopolymers between 11 bp and 20 bp, that are difficult to resolve with the ultima pipeline. For SNVs, in homopolymers between 7 bp and 11 bp the ultima pipeline (precision: 0.955, recall: 0.903) performs slightly worse than the illumina pipeline (precision: 0.994 recall: 0.995). In homopolymers between 12 bp and 20 bp, the performance of ultima further drops (precision: 0.851, recall: 0.727), while illumina is less affected (precision: 0.974, recall: 0.964). The difference in performance between ultima and illumina is much larger for Indels, both for homopolymers between 7 and 11 bp, where ultima (precision: 0.859, recall: 0.822) performs worse than illumina (precision: 0.99, recall: 0.979), but especially for homopolymers between 12 and 20 bp, where the performance of ultima (precision: 0.49, recall: 0.347) further drops with respect to illumina (precision: 0.982, recall: 0.973). Figure 4 shows the top 10 genes in the CMRG benchmark for difference in true positives between ultima and illumina (Δ*TP*), and difference in false positives between ultima and illumina (Δ*FP*).

## Conclusion

In this study we evaluated the small variant calling performance of the Ultima Genomics UG 100™ sequencing platform in comparison with the Illumina NovaSeq 6000 System platform using four genomes of the GIAB Consortium: HG001, HG002, HG003 and HG004. SNV and Indel calling performance was evaluated on the basis of precision and recall computed with hap.py according to the GA4GH best practices in small variant callset benchmarking. In particular, results were stratified using the homopolymer tracks of the GA4GH stratification files, which include homopolymer regions longer than 11 bp that are excluded from the High Confidence Regions bed file (UG HCR), a subset of the NIST v4.2.1 benchmark confident regions that Ultima Genomics uses to benchmark the UG 100™ platform.

Our analysis shows how small variants identified with the ultima pipeline have lower precision and recall with respect to the illumina pipeline (Fig.1 and Fig.2), especially for Indels in homopolymer regions longer than 11 bp (Fig.2). The latest UG Solaris chemistry offers a considerable improvement in performance, although not yet on par with the NovaSeq 6000 System device.

**Fig. 2.**
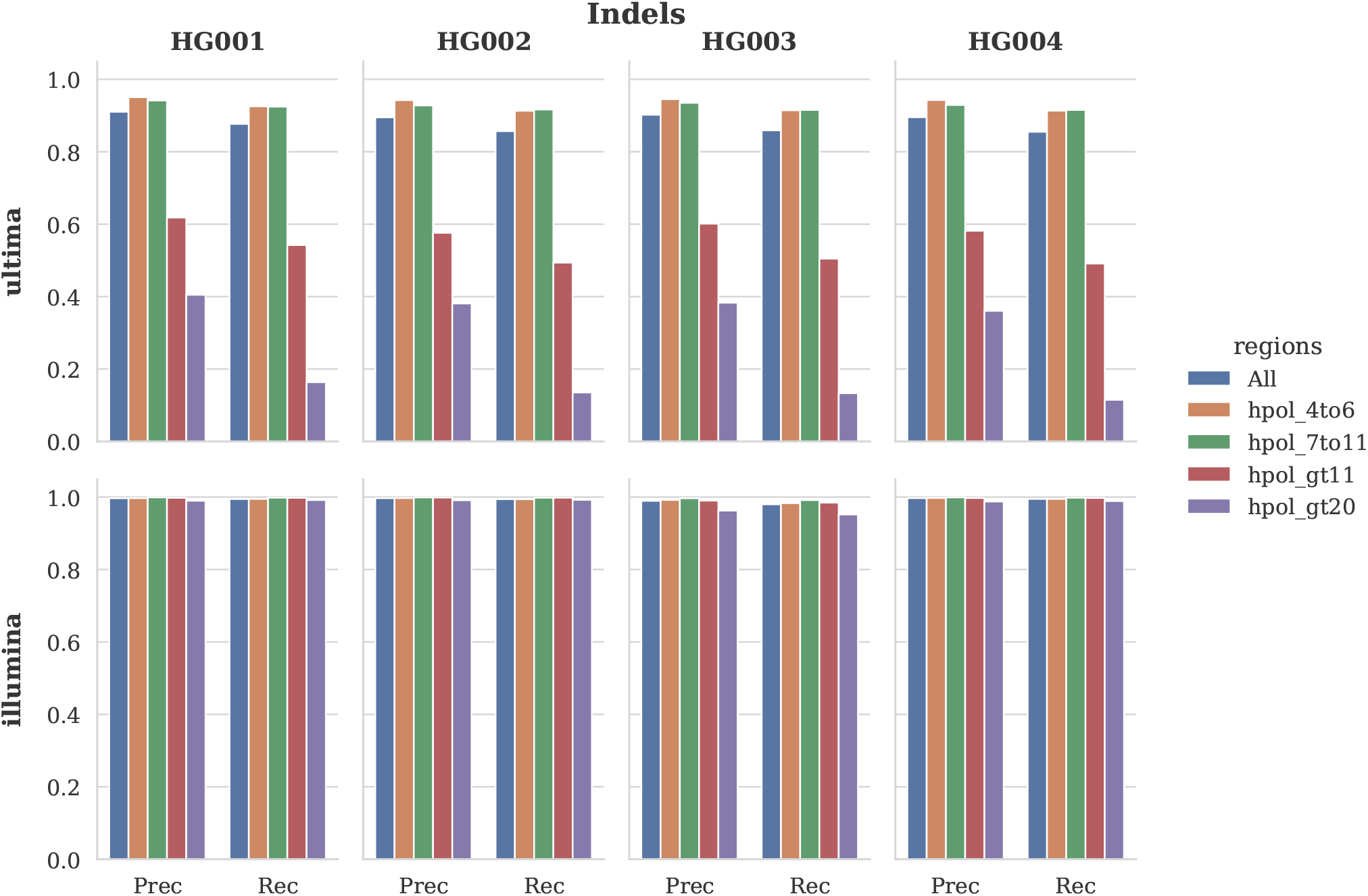
Precision (Prec) and recall (Rec) for Indel calling on the four GIAB samples sequenced with UG 100™ (ultima) and NovaSeq 6000 System (illumina). Genomic regions considered are all benchmark regions (All), homopolymers from 4 to 6 bp (hpol_4to6), homopolymers from 7 to 11 bp (hpol_7to11), homopolymers longer than 11 bp (hpol_gt11) and homopolymers longer than 20 bp (hpol_gt20).

The difference in performance between ultima and illumina is more pronounced in the small variant benchmark of the HG002 Challenging Medically-Relevant Genes (CMRG) (Fig.3). An analysis of small variants in the ClinVar dataset with respect to homopolymer regions shows how the lower performance of ultima in homopolymers longer than 11 bp would affect 69 pathogenic SNVs and 35 pathogenic Indels.

**Fig. 3.**
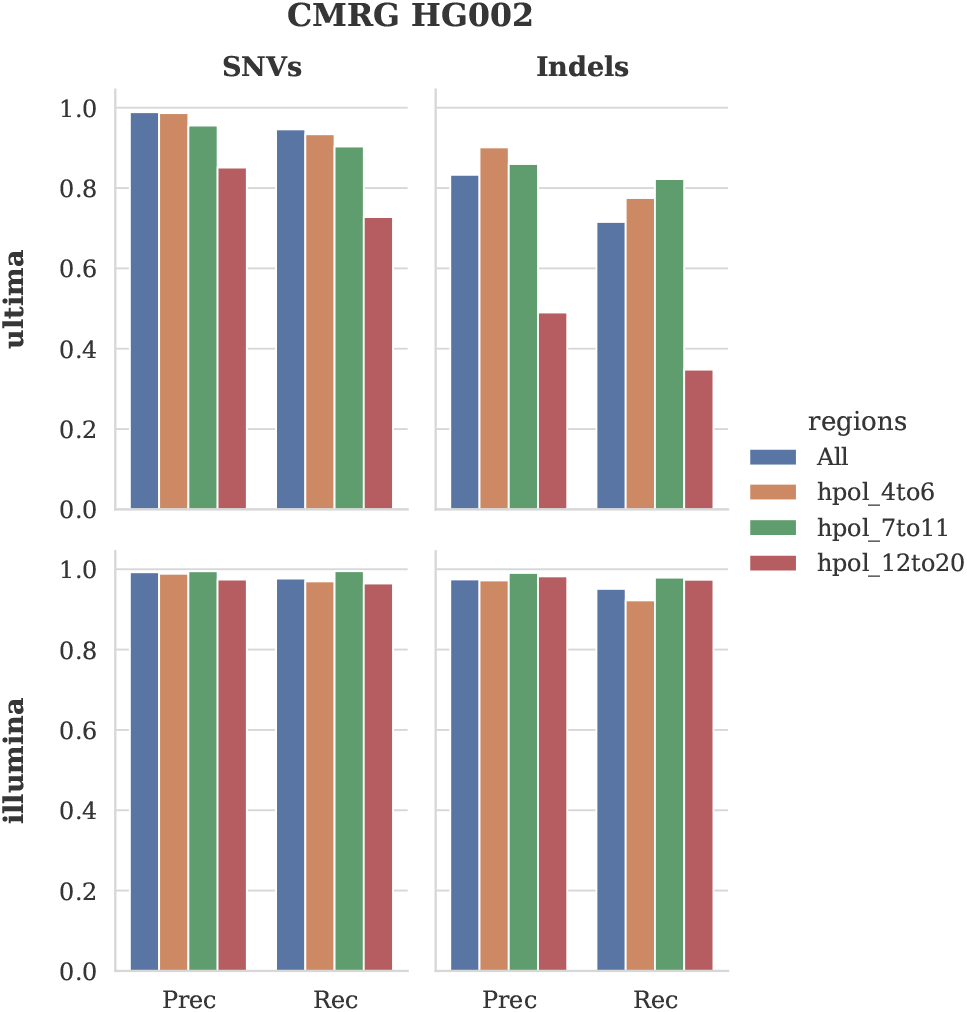
Precision (Prec) and recall (Rec) for SNV and Indel calling on the HG002 GIAB sample sequenced with UG 100™ (ultima) and NovaSeq 6000 System (illumina), considering the benchmark of challenging medically-relevant genes (CMRG).

On October 7^th^, 2025, Illumina published an article [10] with a comparison between the NovaSeq X Series and the Ultima Genomics UG 100™ platforms. In particular, the authors used the publicly available UG 100™ GIAB dataset to show how the UG HCR regions Ultima Genomics currently recommends for the analysis of UG 100™ variants miss ClinVar pathogenic variants in 793 genes. However, this result includes also large variants and it is not stratified by small variant type. The article also shows how the UG 100™ coverage drops in high GC regions, while it remains high for the NovaSeq X Series. In contrast, our study presents a comparison of the UG 100™ and the NovaSeq 6000 System platforms, where the UG 100™ data was generated using the chemistry preceding the Solaris chemistry. SNV and Indel calling performance is stratified using the GA4GH stratification regions for homopolymers, without restricting the analysis to the UG HCR regions. We also show the performance of the two platform on the benchmark set of Challenging Medically-Relevant Genes (CMRG), highlighting the genes for which the number of TP and FP small variants differs between ultima and illumina. For ClinVar variants, we focus on SNVs and Indels stratified by homopolymer regions.

The UG 100™ device is the first sequencing platform to lower the cost of human genome sequencing to 80$. Despite limitations affecting in particular Indel calling in long homopolymer regions, which the latest Solaris chemistry improves upon, this platform offers an unprecedented cost-quality trade-off for increasing the scale of clinical sequencing projects. We hope that our evaluation can guide clinical sequencing centers and clinicians in their choice of the sequencing platform to consider for their clinical studies.

### The GIAB dataset of the pilot study

GIAB samples for the HG001, HG002, HG003 and HG004 genomes were sequenced and analyzed as follows. The Illumina TruSeq DNA PCR-Free protocol was used to prepare the libraries for sequencing with the Illumina NovaSeq 6000 System. DRAGEN v.4.13.3 was used to map Illumina reads on the GRCh38 reference genome (GenBank assembly GCA 000001405.15 no alt analysis set). An additional set of Illumina TruSeq DNA PCR-Free libraries was sent to Ultima Genomics where it was adapted to for sequencing with the UG 100™ platform. The UG pipeline was used to align the reads on the GRCh38 reference genome. The alignment files in CRAM format were downsampled to 30x coverage with *samtools view* to allow comparison with the Illumina data. Small variants were called using the UG VariantCalling workflow that includes the UG DeepVariant version (Efficient DeepVariant). Illumina hap.py v0.3.15 [11] with the RTG Tools 3.13 vcfeval engine (additional parameters: -t ga4gh –pass-only –fixchr) was used to calculate precision and recall on the NIST v.4.2.1 benchmark and on the HG002 CMRG benchmark v1.00, following the best practices for benchmarking germline small variant calls [11]. hap.py results were stratified using the homopolymer tracks (*GRCh38_SimpleRepeat_homopolymer_[hl]_slop5*.*bed*.*gz* : with *hl* as either 4to6, 7to11, ge12, ge21) of the GA4GH genome stratifications version v3.6 [3].

### ClinVar overlap analysis

The ClinVar dataset (clinvar 20251123) was downloaded from the ClinVar website [12]. Only ClinVar SNVs and Indels with a single value for the INFO CLNSIG field equal to either Benign, Likely_benign, Uncertain_significance, Likely pathogenic or Pathogenic, were considered. R v4.4.3 and Bioconductor v3.20 [7], with the packages GenomicRanges v1.58.0 [13], VariantAnnotation v1.52.0 [14] and the tidyverse (readr v2.1.5 and dplyr v1.1.4) [23] R packages, were used to compute the overlap of the ClinVar small variant with respect to the homopolymer tracks, with the ranges of homopolymer lengths being either from 4 to 6 bp (hpol_4to6), from 7 to 11 bp (hpol_7to11), longer than 11 bp (hpol_gt11) or longer than 20 bp (hpol_gt20), of the GA4GH genome stratifications version v3.6 [3]. Figure 4 was created using the Python packages pandas v2.3.3 [15], matplotlib v3.10.7 [8] and adjustText v1.3.0 [4].

Figures 1, 2, 3 and 5 were created with the Python package seaborn v0.13.2 [21].

**Fig. 4.**
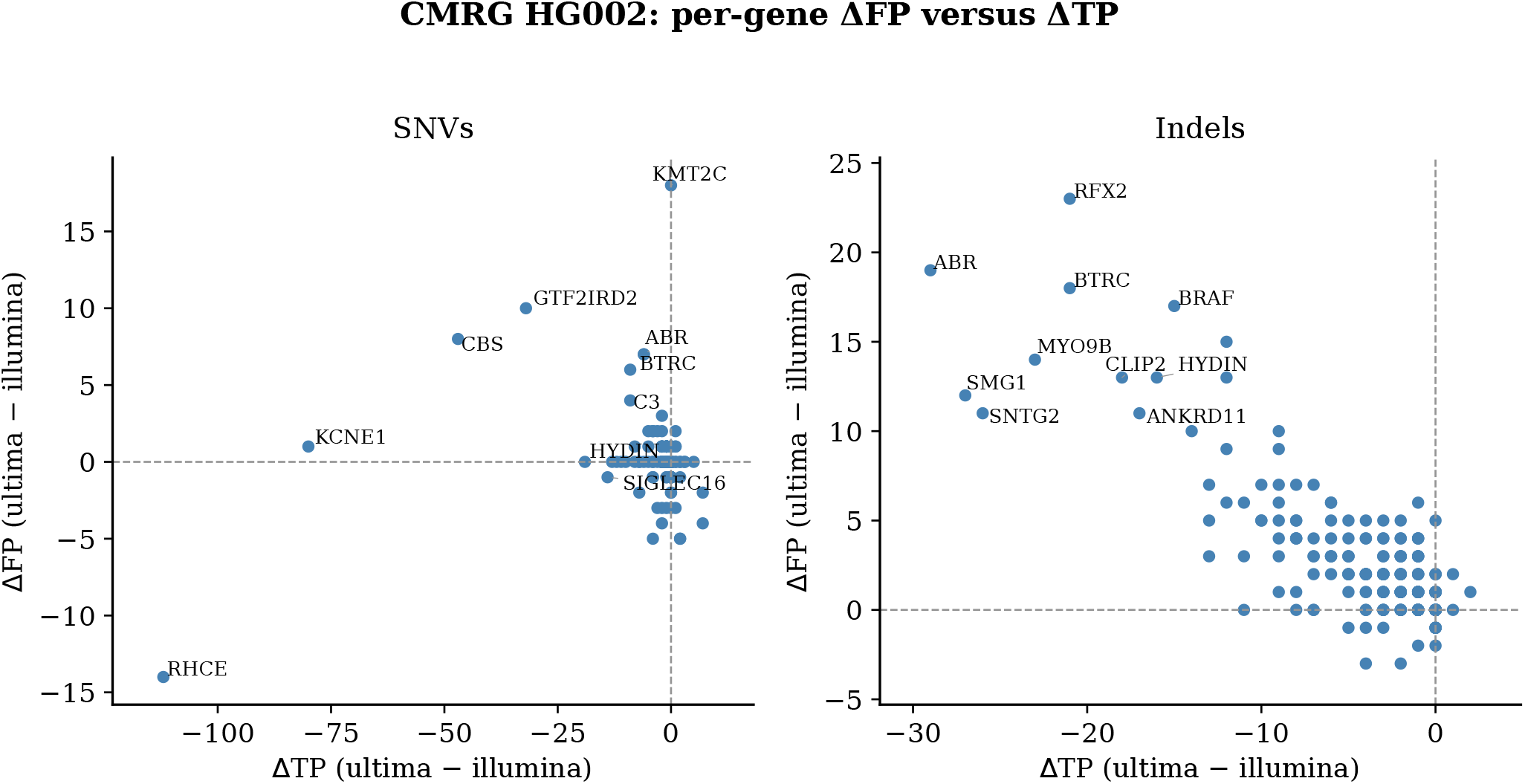
Difference between ultima and illumina FPs (ΔFP) versus TPs (ΔTP) SNVs and Indels for the HG002 CMRG benchmark evaluated with hap.py. The top 10 genes based on (|Δ*TP* | + |Δ*FP* |) are shown.

**Fig. 5.**
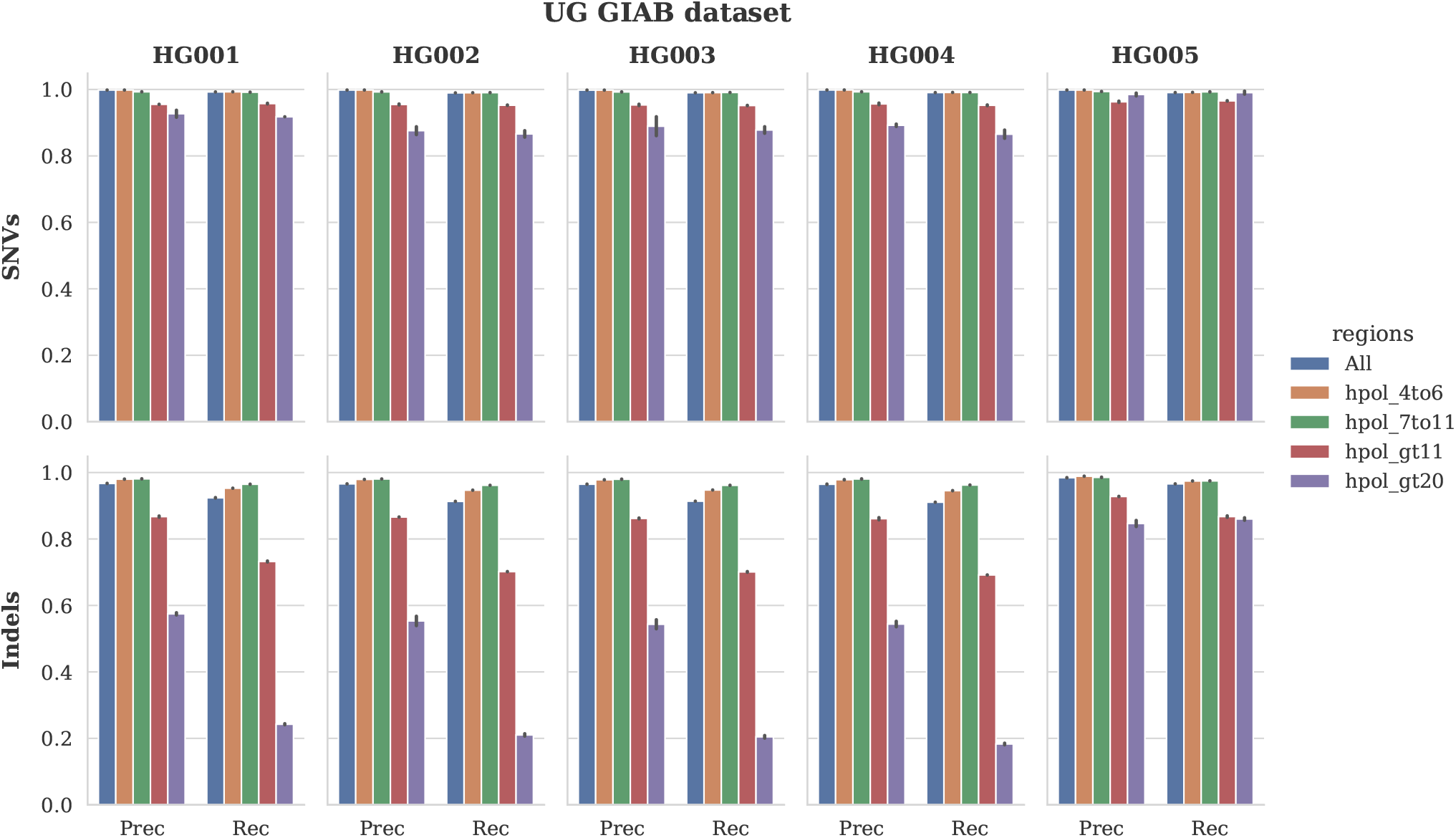
Precision (Prec) and recall (Rec) for SNVs and Indels calling on the five GIAB samples sequenced with UG 100™ (ultima) by Ultima Genomics. Mean and standard deviation (error bars) of two replicates are shown. Genomic regions considered are all benchmark regions (All), homopolymers from 4 to 6 bp (hpol_4to6), homopolymers from 7 to 11 bp (hpol_7to11), homopolymers longer than 11 bp (hpol_gt11) and homopolymers longer than 20 bp (hpol_gt20).

### The Ultima Genomics GIAB dataset

We downloaded the small variant callsets in VCF format for the publicly available GIAB dataset of Ultima Genomics [5]. This dataset includes five genomes: all four genomes considered in our study (HG001, HG002, HG003 and HG004), and the son of the trio from Chinese ancestry sequenced in the Personal Genome Project (NA24631, NIST ID: HG005) [24]. The ultima pipeline has better performance for HG005 in homopolymer regions. Understanding the reason of this difference in performance would require an additional evaluation that is out of the scope of this study.

## Data and code availability

Sequencing data is available on NCBI SRA, Accession: PRJNA1377908 [2]. VCF files have been deposited in Zenodo [18]. The code to reproduce the analysis is available on Zenodo [17].

## Author contributions statement

L.S. conducted the analyses, analysed the results and wrote the manuscript. I.K. conducted the analyses. A.B. prepared the samples for sequencing. I.X. and C.H. supervised the study. All authors reviewed the manuscript.

## Acknowledgments

The authors would like to thank Ultima Genomics for the support in the sequencing with the UG 100™ platform and in the variant calling process using the UG VariantCalling pipeline. The authors would also like to acknowledge all the members of the Health 2030 Genome Center, in particular Sten Ilmjärv for the IT support, Arkadiy Shevrikuko for the data management support and Henri Pegeot for the support in the sample handling.

